# YidC from *Escherichia coli* forms an ion-conducting pore upon activation by ribosomes

**DOI:** 10.1101/2023.05.10.540180

**Authors:** Denis G. Knyazev, Lukas Winter, Andreas Vogt, Sandra Posch, Christine Siligan, Nikolaus Goessweiner-Mohr, Nora Hagleitner-Ertugrul, Hans-Georg Koch, Peter Pohl

**Affiliations:** Institute of Biophysics, Johannes Kepler University Linz, Gruberstrasse 40, A-4020 Linz, Austria; Institut für Biochemie und Molekularbiologie, ZBMZ, Faculty of Medicine, 79104 Freiburg, Germany; Spemann-Graduate School of Biology and Medicine (SGBM), 79104 Freiburg, Germany; Faculty of Biology, Albert-Ludwigs-Universität Freiburg, 79104 Freiburg, Germany

## Abstract

The universally conserved protein YidC aids the insertion and folding of transmembrane polypeptides independently or as a part of the SecYEG translocon complex. In the former scenario, the lipid-exposed YidC surface equipped with a highly conserved positively charged arginine is thought to facilitate membrane insertion of the nascent chain by providing a countercharge for the acidic residues at the polypeptide’s N-terminal region. Here we show that the purified and reconstituted *E. coli* YidC forms an ion-conducting transmembrane pore upon binding a ribosome or ribosome-nascent chain complex (RNC). This pore is closed in the absence of ribosomes. As this pore is not visible in the published monomeric YidC structure, we used AlphaFold to construct the model of a parallel YidC dimer. Experimental evidence for a dimeric assembly comes from our BN-PAGE analysis of native vesicles, fluorescence correlation spectroscopy studies, and single-molecule fluorescence microscopy. In the dimeric model, the conserved positively charged arginine and many residues interacting with nascent chains point into the putative pore. This result suggests the possibility of an alternative mode of YidC-assisted membrane protein insertion.

## Introduction

Protein insertion into the cytoplasmic membrane is an essential process in bacterial physiology. YidC is a member of the Oxa1 family of protein insertases (Vogtle et al., 2022) and is involved in inserting proteins into the bacterial cytoplasmic membrane. Multiple pieces of evidence demonstrate that YidC can act in two modes: independently (Samuelson et al., 2001; Yi et al., 2003; Welte et al., 2012) and assisting the major bacterial protein conducting channel SecYEG (Scotti et al., 2000; Beck et al., 2001; Houben et al., 2002; Zhu et al., 2013). Two partially redundant membrane protein integration systems, YidC and SecY, may be required because the number of SecYEG complexes in the bacterial membrane is relatively small (Kudva et al., 2013). A second insertion site for less complex membrane proteins would ease the transport load of the SecYEG translocon. In support of this, the SRP pathway can deliver some multi-spanning membrane proteins to either SecYEG or YidC, suggesting that some membrane proteins’ insertion pathway is promiscuous (Welte et al., 2012).

In the SecYEG assisting mode, YidC is a part of the SecYEG pore (Sachelaru et al., 2017), where it seems to exist as a monomer that binds to the lateral gate of SecYEG (Sachelaru et al., 2013), most probably via the intercalation of the N-terminal transmembrane domain (Petriman et al., 2018; Nass et al., 2022). In this mode, YidC is suggested to enhance the release of transmembrane helices from the SecY channel, facilitate their subsequent folding (Beck et al., 2001; Houben et al., 2002; Zhu et al., 2013), and even promote the assembly of multi-subunit membrane protein complexes (Wagner et al., 2008). Whether YidC can also form pores for protein insertion when acting independently of the SecYEG translocon is unknown. Crystallographic data suggest that the functional YidC unit is a monomer, as was demonstrated for the truncated and full versions of YidC from *Thermotoga maritima* in detergent micelles (TmYidC) (Xin et al., 2018; Nass et al., 2022), for *Bacillus halodurans* YidC2 (BhYidC) (Kumazaki et al., 2014a), and for *E. coli* YidC (EcYidC) (Kumazaki et al., 2014b), in the lipidic cubic phase. The insertion mechanism inspired by the structural data includes sliding a substrate’s hydrophilic and negatively charged N-terminus through the hydrophobic surface of YidC’s water-filled groove, which contains a conserved arginine residue (R366 in EcYidC) (Chen et al., 2017; Chen et al., 2022).

Exposing a charge to the hydrophobic membrane interior is a rare feature for a transmembrane protein. The associated energetic penalties for (i) ion dehydration (Born energy) and (ii) placing a positive charge into an environment already possessing a positive potential (membrane dipole potential) (Pohl et al., 1998), often force the charged side chains to point away from the hydrophobic lipid core. The voltage sensor of voltage-gated channels may serve as an example (Ahern and Horn, 2004; Chanda et al., 2005). Likewise, establishing water-filled groves is energetically favorable inside the protein – like in potassium or sodium channels (Roux and MacKinnon, 1999) – much less so at the protein-lipid interface.

It is also unclear how YidC’s hydrophylic grove ensures specificity for polypeptides and prevents other small molecules from permeating the membrane at the protein lipid interface. What stops other ions from being pulled into the water-filled grove by the positive charge in its center? How does YidC prevent the membrane barrier to small molecules (Hannesschlaeger et al., 2019) from being compromised?

Polypeptide insertion into the membrane by translocases usually occurs from inside a water-filled pore (Smalinskaite et al., 2022). Recent bioinformatic analyses indicated that the SecY protein channel originated from a dimeric YidC-like protein (Lewis and Hegde, 2021), which suggests that YidC might also function as a pore-forming dimer. The EcYidC contains a large 320 amino acid-long periplasmic loop (P1) between the first two membrane helices, which might facilitate dimerization. This function is supported by the crystal structure of the periplasmic domain of EcYidC, which shows a dimer (Oliver and Paetzel, 2008; Ravaud et al., 2008), and by experiments on native *E. coli* membranes (Boy and Koch, 2009). The dimerization interface on the bacterial YidC may be on TM3 (Kohler et al., 2009) or TM5 (Lewis and Hegde, 2021). For other Oxa1 family members, such as the eukaryotic EMC3 and GET1, dimerization depends on TM5 (McDowell et al., 2020; O’Donnell et al., 2020; Pleiner et al., 2020).

Reconstituted planar lipid bilayers proved helpful in characterizing SecYEG’s pore and the elements required to maintain the membrane barrier to small molecules (Saparov et al., 2007). We now used the same model to test the possibility of pore formation by YidC. We observed channel activity in YidC-containing bilayers in the presence of ribosomes or ribosome-nascent chains (RNCs) of the natural YidC substrate FoC. We also assessed the oligomeric state of the reconstituted YidC in vesicles, IMVs, and planar bilayers with fluorescence correlation spectroscopy (FCS), BN-PAGE, and single-molecule microscopy. Based on a new structural model generated by AlphaFold, we conclude that the YidC dimer may form an ion-conductive pore lined by residues known to interact with nascent chains.

## Results

### Purified and reconstituted YidC displays ion channel activity in the presence of its substrate FoC or empty ribosomes

Liposomes with reconstituted YidC were fused to a pre-formed lipid planar bilayer in the presence of FoC-RNCs, consisting of the transmembrane domain of FoC (residues 1-46), followed by an HA-tag and a TnaC stalling sequence (Wickles et al., 2014)). Vesicles were added from the hyper-osmotic side. The vesicle fusion was triggered by an osmotic gradient across the planar bilayer, which caused a destabilization of the contact area between the proteoliposomes and the planar bilayers. The destabilizing effect is due to osmotic water flux directed from the vesicle towards the hypotonic side (Woodbury and Hall, 1988). Analysis of the single-channel recordings (Fig. 1A) revealed a single-channel conductance of 459 ± 20 pS. According to previous reports, SecYEG-RNC complexes and SecYEG-YidC-RNC complexes exhibit similar conductance values (± 15%). The observation suggests comparable pore sizes. The size was deemed necessary for the two earlier reported pores (SecYEG, SecYEG-YidC) to accommodate the substrate.

**Fig. 1.**
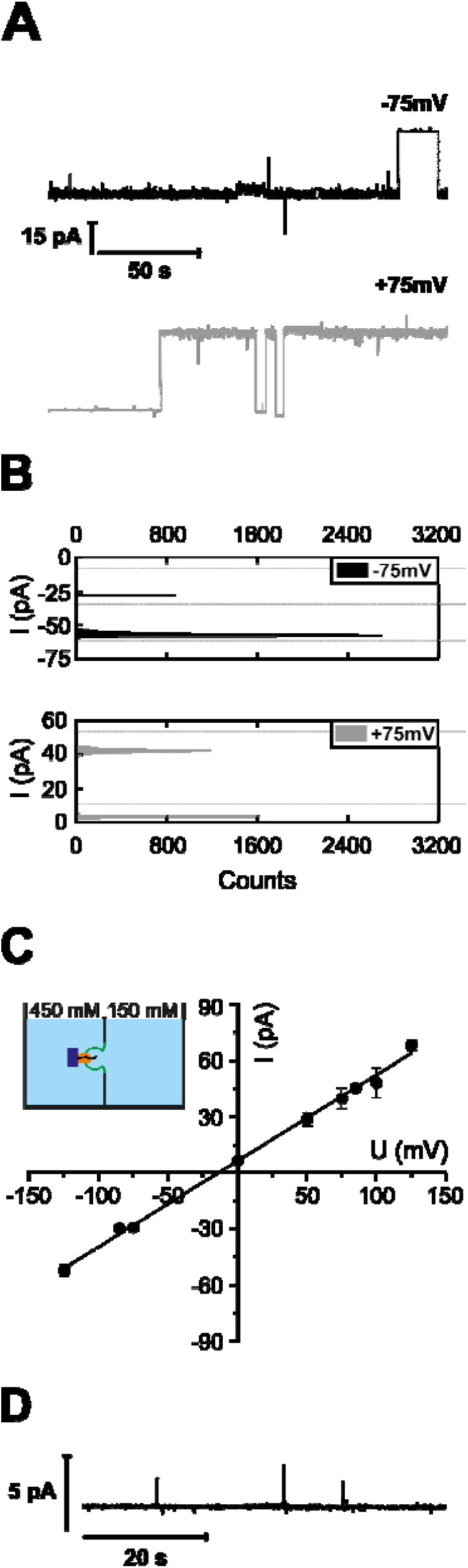
YidC exhibits channel activity after FoC-RNC binding. **(A)** Current traces show single channel openings and closings recorded at different voltages. The YidC-RNC complex was reconstituted into the planar bilayer by vesicle fusion assay conducted in asymmetric salt conditions: 450 mM KCl in the *cis* compartment, 150 mM KCl in the *trans* compartment, and 50 mM K-HEPES, pH 7.5 in both compartments. YidC-proteoliposomes were added to the chamber with pre-formed lipid bilayer from the hypertonic side. The open states are noisier than the closed states due to the frequent flickering of the pore. **(B)** Current histograms corresponding to each trace in (A). The distance between each two neighbor peaks on the histogram corresponds to the current through a single channel. **(C)** The current-voltage plot of the YidC-RNC complex was obtained from single channel amplitudes at different voltages (dots) from traces like in (B). The linear fit yielded the single channel conductivity g = 459 ± 20 pS and a reversal potential of -9 mV resulting from the slight asymmetry in the conductance for cations and anions. A colored scheme shows the chamber with two compartments filled with the solution of the indicated ionic strength. The compartments are separated by the lipid bilayer (green), containing YidC (orange) and bound RNC (ribosome as a purple rectangle with a nascent chain as a black line). **(D)** Negative control: lack of channel activity in the absence of FoC-RNC.

The YidC channel exhibits a reversal potential of -9 mV (Fig. 1B). According to the Goldman-Hodgkin-Katz equation, it indicates a less than twofold preference for anions over cations (see *Calculation of channel ion selectivity* in *Methods*). We previously observed such preference for the SecYEG translocon (Knyazev et al., 2014). The modest anion selectivity makes only a very minor contribution to maintaining the proton motive force during protein translocation.

The addition of (i) FoC-RNCs to protein-free bilayers or (ii) YidC-vesicles to the hypertonic side without FoC-RNC induced no channel activity (Fig. 1D). Hence, the observed channel activity (Fig. 1A-C) was due to the complex of YidC and FoC-RNC.

Even though the ability of YidC to form conductive pores in the presence of the SecYEG translocon and RNCs was reported previously (Sachelaru et al., 2017), we are unaware of any reports about water or ion conductance of substrate-activated YidC in the absence of SecY. Yet, the protein family member Oxa1 forms pores capable of accommodating a translocating protein segment (Kruger et al., 2012).

Since SecYEG channels open upon ribosome binding (Knyazev et al., 2013; Knyazev et al., 2020), we tested whether YidC possesses the same capability. Instead of FoC-RNC (Fig. 1), we added non-translating ribosomes and YidC-containing proteoliposomes to the hypertonic side. Subjecting the induced channel activity (Fig. 2A) to a histogram analysis (Fig. 2B) revealed a unitary channel conductivity g = 436 ± 20 pS, (Fig. 2C) close to 459 ± 20 pS of the FoC-RNC-YidC complex (Fig. 1). However, the RNC concentration needed for observing channel activity was two to three orders of magnitude lower than that of non-translating ribosomes. The reversal potential U_rev_ = -5 mV was even smaller than in the presence of an RNC (Fig. 1), indicating an even smaller preference for anions in the presence of empty ribosomes as compared to FoC-RNC.

**Fig. 2.**
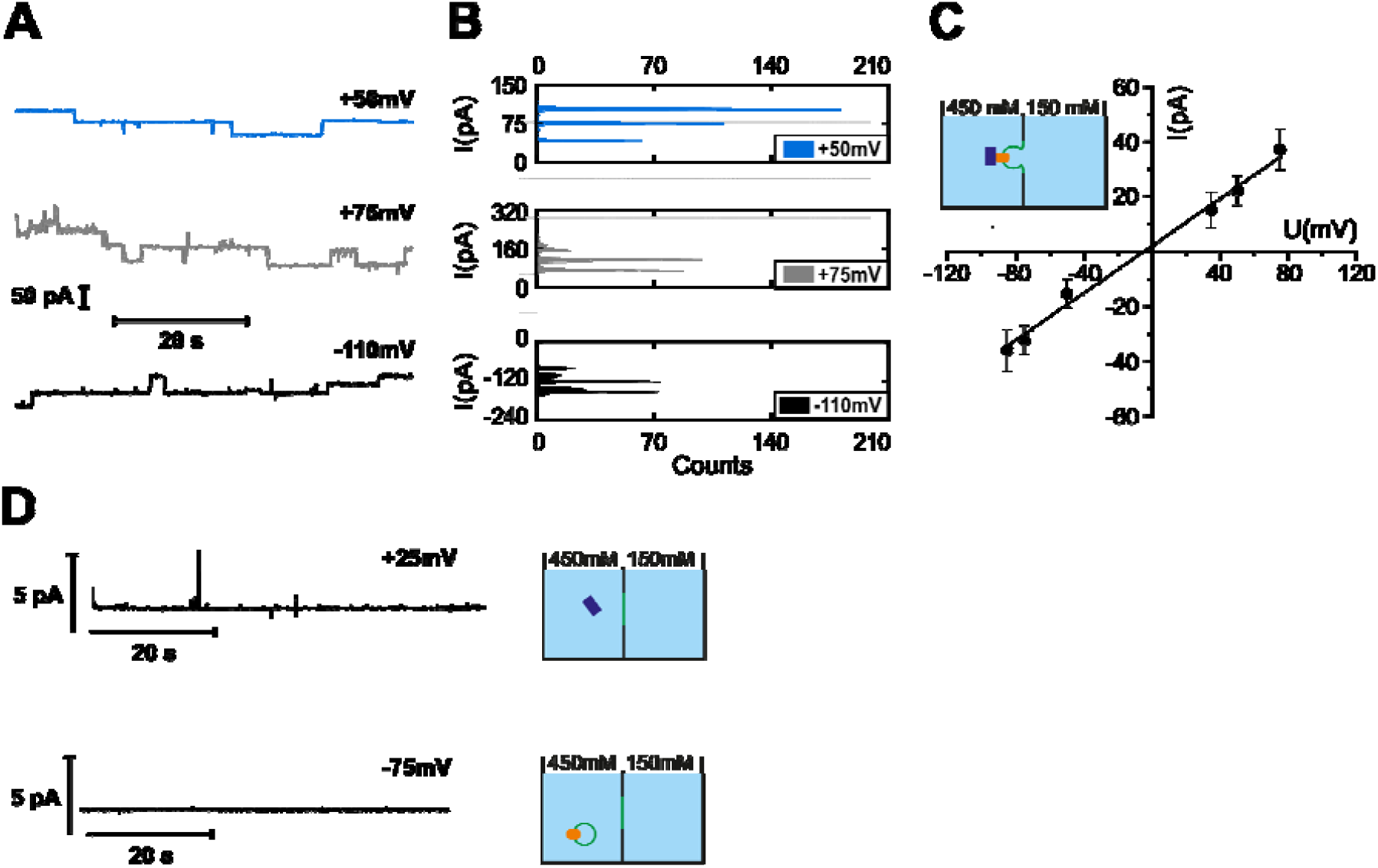
YidC exhibits channel activity after ribosome binding. **(A)** Current traces show single channel openings and closings recorded at different voltages. **(B)** Current histograms corresponding to each trace in (A). The distance between each two neighbor peaks on the histogram corresponds to the current through a single channel. **(C)** The current-voltage plot of the YidC-ribosome complex was obtained from single channel amplitudes at different voltages (dots) from traces like in (B). The linear fit yielded the single channel conductivity g = 436 ± 20 pS. The reversal potential of -5 mV indicates the lower asymmetry for cations and anions’ permeabilities, compared to Fig.1. **(D)** Negative controls: lack of channel activity without ribosomes (lower trace) or YidC (upper trace). The YidC-ribosome complex was reconstituted into the planar bilayer by vesicle fusion, as in Fig. 1.

Adding the YidC-vesicles in the absence of ribosomes (Fig.2D, lower trace) or ribosomes in the absence of YidC vesicles (Fig. 2D, upper trace) did not lead to channel activity. The former proves that YidC is electrically silent without a binding partner, and the latter proves that purified ribosomes did not contain any channel-forming contaminants.

The only other channel which ribosomes could activate is SecYEG. Its co-purification together with YidC is unlikely, as confirmed by Western Blot analysis (SFig. 1). Records with a high number of YidC channels in a single membrane (SFig. 2) also prove that the channel-forming entity cannot be a minor contaminant in the purified sample.

The ligand-free YidC remains electrically silent even when exposed to transmembrane voltage (SFig. 3). Instead of using vesicle fusion to insert the protein into the planar bilayer, we folded solvent-depleted planar bilayers from the monolayers that form on top of proteoliposme suspensions (Saparov et al., 2007). These monolayers also contain membrane proteins (Schurholz and Schindler, 1991). The thus reconstituted YidC was functionally intact, as demonstrated by ribosome-induced channel formation (SFig.3).

Next, we tested whether signal peptides may activate YidC without ribosomes. The membrane insertion of phage proteins (PF3, M13) that are too short to be co-translationally targeted to YidC (Samuelson et al., 2000; Samuelson et al., 2001) supports the hypothesis. Yet the signal peptide of subunit A of cytochrome o oxidase did not activate YidC. We did not observe channel activity unless we added ribosomes. Noteably, cyoA insertion requires both SecY and YidC (du Plessis et al., 2006; van Bloois et al., 2006).

### YidC with C-terminal deletion retains ribosome binding activity

The C-terminus is part of the ribosome-binding site in *E. coli* YidC (Welte et al., 2012). Similarily, the much longer C-terminus of the yeast YidC homolog Oxa1 is also known to be involved in ribosome binding (Szyrach et al., 2003). For electrophysiological experiments with the C-terminal deletion version of YidC (YidC*Δ*C), we folded the bilayers from YidC containing monolayers. The addition of purified empty *E*.*coli* ribosomes led to the formation of ion-conducting channels of conductance g = 392 ± 30 pS (Fig. 3A, B), which is close to the one observed for the complex of empty ribosomes with the full-sized YidC. In line with our result is the observation that deleting the C-terminus reduces ribosome affinity by only approx. 75% (Kedrov et al., 2013). The residual affinity is large enough to result in a significant binding probability.

**Figure 3.**
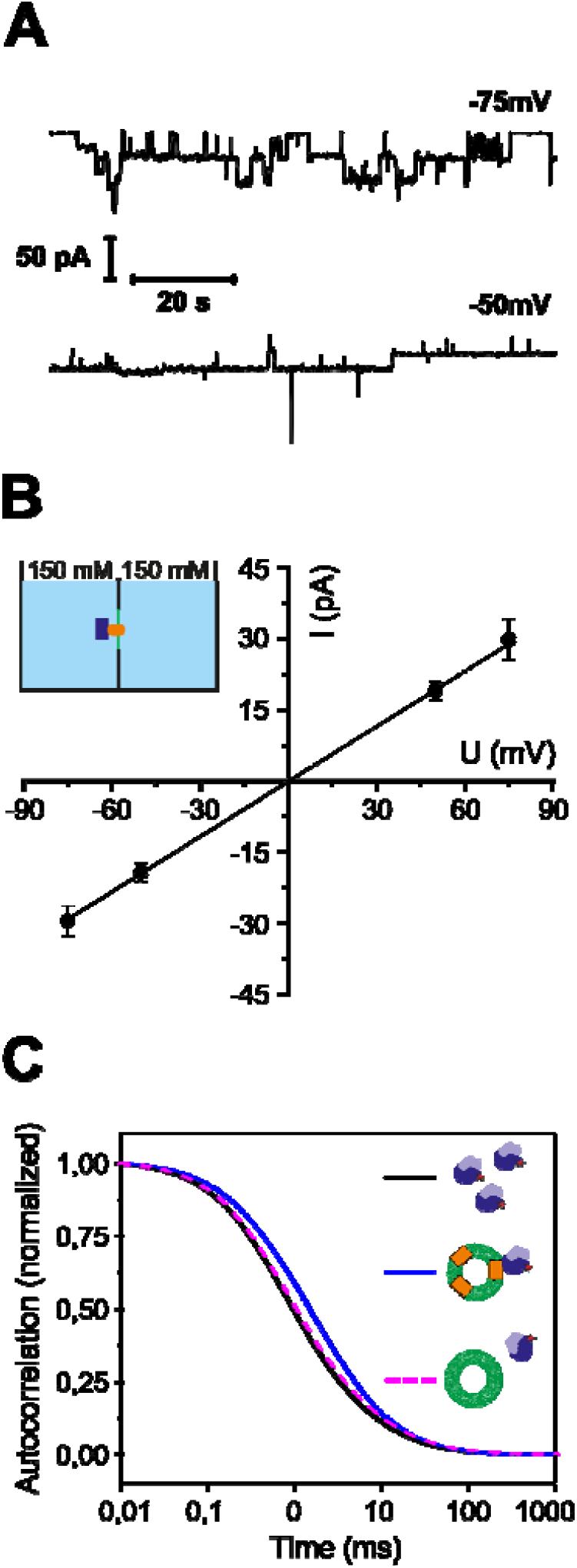
Ribosomes activate YidCΔC. **(A)** Channel activity of the planar bilayers with the reconstituted YidCΔC after the addition of ribosomes. The experiments are analogous to the one in SI Fig. 1. **(B)** Single channel amplitudes (dots) and Current-Voltage characteristic of YidCΔC-ribosome complex (line) at different voltages obtained from traces like in (B). Single channel conductivity g = 392 ± 30 pS. **(C)** The binding of ribosomes to YidCΔC-vesicles observed with Fluorescence Correlation Spectroscopy. Autocorrelation curves of labeled ribosomes in solution (black), of ribosomes mixed with YidCΔC proteoliposomes (blue), and ribosomes mixed with empty vesicles containing lipid label (pink dashed line). The ribosomes and vesicles have comparable diffusion time across the confocal volume, whereas their complex diffuses slower. No binding could be observed when ribosomes were mixed with empty vesicles. Experiments were conducted in 150 mM KCl and 50 mM K-HEPES, pH 7.5.

To confirm this observation, we non-specifically labeled the His-tagged ribosomes with an ATTO-633-NHS ester (Blanchard et al., 2004) and measured their diffusion by FCS in solution in the presence of empty vesicles or YidC proteoliposomes. Upon proteoliposome addition, 40% of the ribosomes showed an increased residence time (Fig. 3C), which indicates the binding of the 70S subuint to the YidC-vesicles.

Our data show that ribosomes can interact with YidC without its C-terminus, supporting previous studies (Kedrov et al., 2013). The His-tag introduced into all our YidC constructs for affinity purification might facilitate the residual binding of ribosomes to YidCΔC. Yet such His-tag mediated binding would not lead to channel opening. Likely, this binding is due to additional positively charged residues within the extended cytosolic loop between TM2 and TM3 (Geng et al., 2015). This may be necessary because the C-terminus of *E. coli* YidC is much shorter than the typical C-terminal ribosome-binding sites in other bacterial species like *Streptococcus mutans* or *Rhodopirellula baltica* (Dong et al., 2008; Seitl et al., 2014).

### Structural YidC models

The crystal structure shows YidC as a monomer (Kumazaki et al., 2014a). Five transmembrane helices form a positively charged hydrophilic groove facing the lipid bilayer. They are arranged in a fashion that does not leave room for a membrane-spanning pore. A subsequent structural model of YidC based on evolutionary co-variation analysis, supplemented by a cryo-electron microscopy reconstruction of a translating YidC-ribosome complex carrying the YidC substrate FoC (Wickles et al., 2014) is in line with this conclusion. The model features a single YidC copy bound to the ribosomal tunnel exit.

The apparent contrast between the pore-free structural models of YidC and our observation of YidC’s channel activity prompted us to interrogate AlphaFold for potential dimer configurations. First, we discarded the resulting antiparallel dimer models since the transmembrane transfer of huge loops is improbable. Moreover, previous research exclusively suggests the existence of parallel dimers (Koch et al., 2002; Serek et al., 2004; Lotz et al., 2008). Second, we selected the parallel dimer with a visible hydrophobic belt, characteristic of membrane proteins (Fig. 4A). The resulting model (Fig. 4B, C) shows a structure that may expand to form a pore in its center when ligand-activated, e.g., by ribosome binding. Both the N-terminal helix, separated from the rest of the protein by a large periplasmic domain, and the C-terminal helix, exposed to the cytosol and thus offering a potential ribosome binding site, may move outwards. The thus formed pore would accommodate (i) many of the residues known to interact with nascent chains (Klenner and Kuhn, 2012) and (ii) the conserved arginine previously described as essential for YidC functioning (Kumazaki et al., 2014a).

**Fig. 4.**
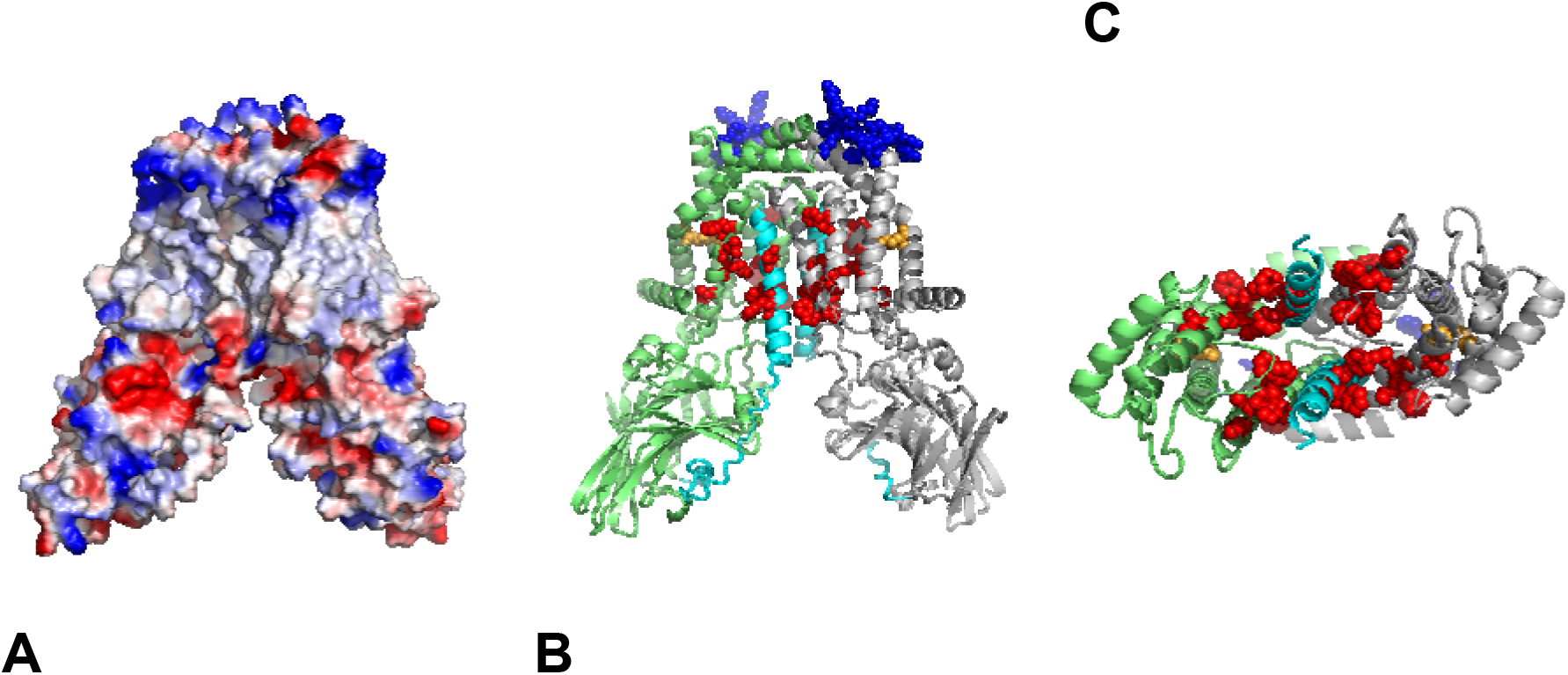
Exploring a potential YidC dimer fold modeled by AlphaFold. (A) Surface charge representation of the parallel dimer model. The two closely interacting alpha-helical domains form a consistent hydrophobic belt. (B) Cartoon representation of the parallel dimer model (top: cytosol, bottom: periplasm). The C-terminal amino acids (blue spheres) are exposed and thus available for potential interaction with a ribosome. The highly flexible N-terminus (cyan) was modeled to participate in the parallel helix formation and might contribute to the hydrophobic belt and substrate translocation or gating. Amino acids previously described as involved in substrate interaction are highlighted with red (Klenner and Kuhn, 2012) and orange (R366) (Kumazaki et al., 2014a) spheres, respectively. (C) Model viewed from below and with the beta-domains removed. Notably, most of the amino acids described as potential interactors with YidC substrates are grouped in the center of the parallel complex, many of them even facing to the inside. The model was analyzed and visualized with PyMOL (Schrödinger).

### Stoichiometry of reconstituted YidC

Next, we determined the oligomeric state of YidC in proteoliposomes. Therefore, we fluorescently labeled YidC with Atto-488-maleimide and used FCS to count the number of YidC molecules per liposome as described previously (Knyazev et al., 2013). In short, fluorescence correlation spectroscopy (FCS) served to determine the number of proteoliposomes per focal volume. Dissolving them with a mild detergent (3 % octyl glucoside, OG) increased the number of fluorescent particles three-fold. Each fluorescent particle corresponded to a lipid-detergent-protein micelle (Fig. 5A). Subsequent addition of a harsh detergent (1.1 % sodium dodecyl sulfate, SDS), doubled the number of particles, indicating that OG-micelles contained an average of one YidC dimer. That is, each liposome harbored about three YidC dimers.

**Figure 5.**
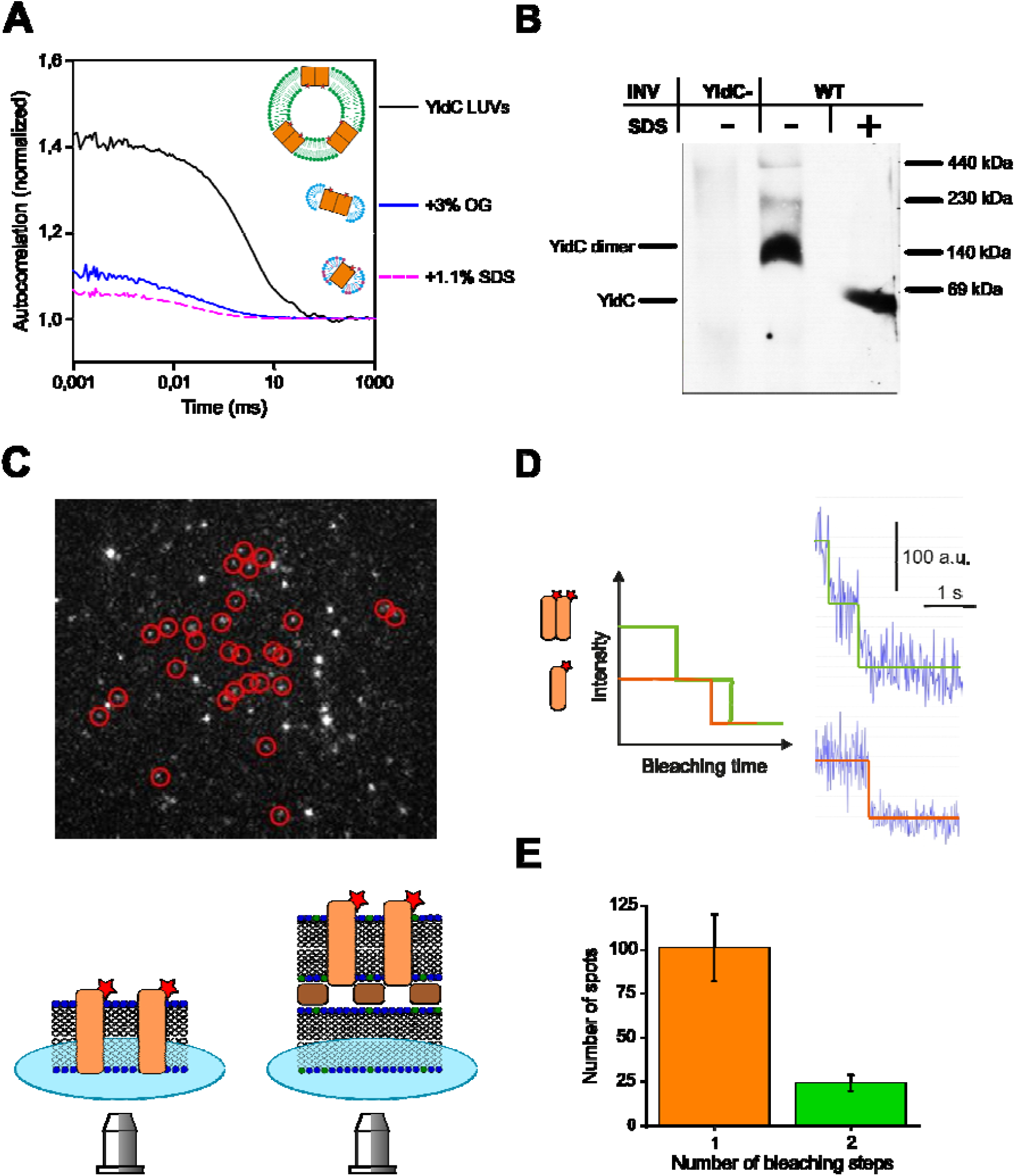
Stoichiometry of YidC. (A) YidC stoichiometry in reconstituted liposomes was obtained with FCS. Vesicles used in electrophysiological experiments were obtained with the same reconstitution procedure. The number per confocal volume of YidC-vesicles, YidC-OG-lipid, and YidC-OG-lipid-SDS micelles was derived from the amplitude A of the corresponding autocorrelation curve as 1/(A-1). In the same order, the number of resulting particles amounted to about 2.5, 7.5, and 15, respectively, meaning that the vesicles contained, on average, three YidC dimers. The color code for cartoon schemes is the same as in Fig. 3, with OG molecules depicted in cyan and SDS molecules in dark red. The red stars indicate the fluorescent label. (B) YidC forms dimers in its native environment. INVs of wild-type *E. coli* cells (wt) or a conditional YidC depletion strain (YidC-) were analyzed by BN-PAGE and, in course of western blotting, decorated with an antibody against YidC. The BN page clearly shows a YidC dimer at 140 kDa (lane 2), which can be dissolved to a monomeric state by SDS treatment before loading the BN-PAGE (lane 3). (C) An exemplary Single Molecule Fluorescence Bleaching experiment. Such experiments were conducted on supported (lower scheme on the left) and suspended (lower scheme on the right) lipid bilayers which were formed from YidC-vesicles mixed with empty vesicles (1:1000). The color code of the schemes is the same as above, with streptavidin crystal shown as brown rectangles, biotinylated lipid shown in green, and cover slide glass in blue. The spots marked by red circles were later analyzed. The brighter spots correspond to aggregates. (D) Spots in the red circles showed one- and two-step bleaching as schematically shown on the left, with exemplary intensity traces on the right. (E) The histogram of the number of bleaching steps generated after analyzing the spots from different experiments shows that 101 spots were bleached in one step and 24 in two steps. That is, approx. 1/5 of the spots corresponded to YidC dimers.

To eliminate the possibility that interactions with the detergent affected YidC’s oligomeric state, we performed blue native PAGE analysis of *E. coli* inverted inner membrane vesicles (IMVs) from the wild-type cells with native level of YidC expression. In the absence of SDS, Western blot analysis showed an abundance of YidC dimers, which agrees with previous data (Boy and Koch, 2009). In contrast, in SDS, we detected only the monomeric form (Fig. 5B). Our results demonstrate a significant imerization propensity of YidC at its cellular concentration.

Finally, we performed single-molecule fluorescence experiments to prove that YidC dimers were present in planar lipid bilayers. Since the fluorescence intensity of single fluorophores observable in free-standing planar bilayers is significantly lower than in bilayers near the objective, we performed the stepwise bleaching experiments of labeled YidC complexes in supported and suspended lipid bilayers. The latter model removes the protein from the support because the protein harboring membrane rests on a layer of streptavidin crystals (Zhu et al., 2016), as schematically shown in Fig. 5C. Membrane proteins are free to diffuse in the valleys between streptavidin pillars. To form supported or suspended bilayers, we used empty vesicles and proteoliposomes in a ratio of 1000:1. We analyzed the bleaching of individual fluorescent spots (Fig. 5C, D). It appeared that, apart from way larger aggregates which showed a gradual decrease of fluorescence during bleaching, there were two major populations of particles. Particles in the first population bleached in one step and those in the second in two. As both types of bilayers, supported and suspended, showed similar results, we pooled together the data from both systems for the monomer-dimer distribution (Fig. 5E). Thus, a fifth of the observed complexes, corresponding to one-third of the total YidC molecules, existed as dimers, with the rest forming monomers.

## Discussion

The insertase YidC mainly facilitates the transfer of single-spanning membrane proteins in a Sec-independent manner. It can also enter the SecYEG complex and the SecDF complex and form complexes with other membrane proteins, such as PpiD and YfgM (Götzke et al., 2014; Sachelaru et al., 2017; Jauss et al., 2019; Oswald et al., 2021; Watkins and Collinson, 2022). YidC’s envisioned polypeptide insertion mechanism does not require a pore – in contrast to the operation mode of translocons (e.g., SecYEG). The insertase merely primes the lipid bilayer at its interface for polypeptide insertion. It is unknown how the membrane barrier to small molecules, like ions, is maintained by the primed bilayer or during polypeptide insertion.

In contrast, by exploiting an elaborate gating mechanism, the SecYEG translocon maintains the barrier to small molecules. Its pore is closed in the resting state (Saparov et al., 2007), opens upon binding of a signal sequence (Simon and Blobel, 1992), other ligands (Knyazev et al., 2013; Knyazev et al., 2014), and responds to membrane potential (Knyazev et al., 2020). Observing a gated YidC pore raises the question of whether it may serve the same purpose. If it were used for protein insertion, the pore would provide the means for maintaining the barrier to small molecules while allowing the passage of large polypeptides. Observing a dimeric configuration that accommodates many residues interacting with the polypeptide (Fig. 4) supports the notion. Additionally, single-channel conductivity indicates that channel diameter is only 20 % smaller than that of SecYEG (Saparov et al., 2007), i.e., the pore would be large enough to accommodate an unfolded polypeptide chain.

The pore-forming ability of YidC is further supported by electrophysiological studies on Oxa1, the YidC homolog in the mitochondrial inner membrane (Kruger et al., 2012). Oxa1 was shown to form a cation-selective channel that responds to mitochondrial export signals. However, the Oxa1 channel appears larger than the YidC channel, which could reflect Oxa1’s propensity to form tetramers in the mitochondrial membrane (Nargang et al., 2002; Reif et al., 2005). Oxa1 is primarily responsible for inserting the hydrophobic core subunits of the respiratory chain complexes (Kulawiak et al., 2013). Thus, Oxa1 has very restricted substrate specificity. This differs from EcYidC, which can insert membrane proteins with varying hydrophobicity and polarity (Soman et al., 2014).

As we show here, the ribosome also interacts with the C-terminal deletion YidC mutant, meaning that another interaction site in addition to the proposed C-terminus (Kohler et al., 2009) is present, for example, loops between helixes 2-3 and 4-5, as was reported in (Wickles et al., 2014; Geng et al., 2015).

Our BN-PAGE and FCS experiments with YidC-vesicles and IMVs showed a predominantly dimeric form both in native *E. coli* membranes and reconstituted proteoliposomes (Fig. 5A, B). In comparison, the single-molecule experiments on the suspended and supported bilayers with lower YidC-to-lipid ratios due to the dilution of YidC required for single-molecule microscopy showed a predominantly monomeric form (Fig. 5C-E). We conclude that YidC can exist in dimeric and monomeric forms in the lipid bilayer, with an equilibrium between the dimeric and monomeric forms.

The structural model built by AlphaFold (Fig. 4) supports the hypothesis that the dimer is the pore-harboring entity. The observation of eight fusion events with an average amplitude of ∼54 pA corroborates the model (SFig. 2). This amplitude indicates the presence of roughly three YidC channels per vesicle. At the same time, FCS revealed the presence of three YidC dimers per vesicle (Fig. 5A), suggesting that only one channel may be formed per dimer. The conclusion is in line with the previously made observation that, in a genetically fused YidC dimer, each monomer can act independently of the other (Spann et al., 2018).

Although compelled by the visualization of pore-lining interaction sites with the nascent chain, the projection that only the dimer facilitates polypeptide insertion may not be valid. A cryo-electron microscopy study puts the nascent chain on the surface of a ribosome-bound YidC monomer (Wickles et al., 2014). The low resolution of the structure may allow other interpretations, yet evidence that the dimer may harbor the nascent chain is thus far missing. Additional work is required to demonstrate the functional significance of the YidC dimer. Moreover, there is a distinct possibility that the dimerization propensity of YidC and pore formation might be important for only a subset of substrates, in particular those that are multi-spanning and can also be inserted by SecY, e.g. TatC.

## Materials and Methods

### Cloning of pTrc99a-YidCΔC and construction of a single-cysteine YidC derivative

pTrc99a-YidC*Δ*C was cloned from pTrc99a-YidC (Welte et al., 2012) by inverse PCR deleting the last 13 amino acids of YidC using the following primer pair:

YidCdeltaCRev: (5’GCTGATTTACCGTGGTCTG3’); YidCdeltaCFwd: (5’-TGATTCGGTGAGTTT TCG-3’). For the labeling of YidC with ATTO488, a single cysteine mutation was introduced by site-directed mutagenesis at position 269 (YidC_D269C_fwd: 5’-CCCGCATAACTGCGGTACCAACAACTTC-3’; YidC_D269C_rev: 5’-GAAGTTGTTGGTACCGCAGTTATGCGGG-3’) in a cysteine free YidC (C423S) generated by site-directed mutagenesis from pTrc99a-YidC (Welte et al., 2012) (YidC_C423S_fwd: (5’-CCGCTGGGCGGCTCCTTCCCGCTG-3’; YidC_C423S_rev: G CAG CGG GAA GGA GCC GCC CAG CGG). The D269C has been previously used for fluorescent labeling of YidC (Kedrov et al. 2013).

### Purification and reconstitution of YidC into vesicles

YidC and YidC*Δ*C were purified from pTrc99a-YidC (Welte et al., 2012) and pTrc99a-YidC*Δ*C expressing BL21 cells using a Ni-NTA FF crude column (GE Healthcare) on an ÄKTA chromatography system. The equilibration/wash buffer contained 50 mM Tris-HCl pH 7.5, 300 mM NaCl, 5 mM MgCl_2_, 20 mM imidazole, 0.03 % DDM (Affymetrix Antrace), 10 % glycerol, and His-tagged proteins were eluted with a linear gradient from 20 to 500 mM imidazole. Dried *E. coli* polar phospholipids (Avanti Polar Lipids) were rehydrated in buffer (50 mM Tea-OAc pH. 7.5, 50 mM DTT) to a final concentration of 100 mg/ml and sonicated. Proteoliposomes were prepared by mixing lipids (final concentrations of 0.1 mg/ml) and 0.85 % (w/v) n-octyl-β-D-glycoside with 1.5 μM purified, and DDM-solubilized proteins and incubating them for 20 min at 4° C. Samples were then dialyzed (Spectrapor membrane tubing, 6-8 kDa) against 50 mM TeaOAc, pH 7.5, 1 mM DTT. The proteoliposomes were pelleted (1 h, 210 000xg) and resuspended in 50 mM TeaOAc, pH7.5, 1 mM DTT to a final protein concentration of 5 μM. YidC-to-lipid ratio was 1:100, m%. The reconstituted YidC is functionally active, as indicated by its ability to insert different membrane proteins like MtlA or TatC (Welte et al., 2012).

### Labeling of YidC

YidC(D269C, C423S) was labeled with ATTO488 on the periplasmic loop before reconstitution. EcYidC’s loop is almost as big as BhYidC’s. Therefore the purified protein was incubated with 100 *μ*M TCEP for 5 min on ice before incubating with a ten-fold molar excess of ATTO-488-maleimide (Sigma-Aldrich) for 2 hours on ice. Excess dye was removed by desalting via PD10 columns (GE Healthcare) before reconstitution.

### BN-PAGE analysis

Purified IMVs (100 μg protein) of wild-type *E. coli* TY0 cells or the conditional YidC depletion strain JS7131 (Urbanus et al., 2002) were dissolved in a buffer containing 50 mM imidazole/HCl, pH 7.0, 5 mM 6-aminocaproic acid, 50 mM NaCl, solubilized with 1 % final concentration n-Dodecyl β-D-maltoside (Roche, Mannheim, Germany) and incubated for 5 min at 25 °C. Non-solubilized material was pelleted by centrifugation for 30 minutes at 45.000 rpm, 4 °C (TLA-45 Beckman rotor). The solubilized proteins were separated on 4–15 % BN-gels and analyzed by immune detection using polyclonal α-YidC antibodies. Dissociation of YidC dimers was induced by treatment with 0.1 % SDS for 10 min at 56 °C before loading on BN-PAGE.

### CyoA leader peptide

We tested the signal peptide of the precursor form of subunit A of the cytochrome o oxidase (sequence: MRLRKYNKSLGWLSLFAGTVLLSG). It was synthesized and purified to 98 % purity by Peptide 2.0 Inc. (Chantilly, USA).

### Reconstitution of YidC into planar bilayers

Planar bilayers were formed in a Teflon chamber with two compartments separated by a Teflon septum. Pure lipid vesicles from E.coli polar lipid extract (Avanti Polar Lipids, Alabaster, AL) were added to the two compartments containing 50 mM K-HEPES and 150 mM KCl (pH=7.5). The final lipid concentration was between 1 and 2 mg/ml. Planar bilayers were then folded by raising the level of these two aqueous solutions over the dividing aperture in a Teflon septum, thereby combining the two lipid monolayers on top (Hannesschlaeger and Pohl, 2018). 30 to 60 minutes of control current recordings ensured the absence of lipid channels.

We used two different approaches for the subsequent protein reconstitution: (i) We lowered the level of the aqueous solutions below the aperture, added YidC containing proteoliposomes, and raised the buffer levels above the aperture after an incubation time of ∼1 hour (Pohl et al., 2001; Saparov et al., 2007). (ii) We added proteoliposomes and ribosomes to one side of the intact membrane. Increasing the osmolarity in that compartment by adding 300 mM KCl resulted in YidC insertion into the planar membrane by vesicle fusion (Woodbury and Hall, 1988). Ribosomes were added to a final concentration of 0.5 – 1 mg/ml. For experiments with ribosome nascent chain complexes, RNCs were added to a final concentration of 0.5-1 μg/ml.

### Single ion channel measurements

Ag/AgCl reference electrodes were immersed into the buffer solutions on both sides of the lipid bilayer. The command electrode of the patch clamp amplifier (model EPC9, HEKA electronics, Germany) was immersed into the cis compartment, and the ground electrode into the trans compartment. The recording filter for the transmembrane current was a 4-pole Bessel with a -3 dB corner frequency of 0.1 kHz. The raw data we obtained were analyzed using the TAC software package (Bruxton Corporation, Seattle, WA). Gaussian filters of 12 Hz were applied to reduce noise.

### Ribosome expression and purification

Tetra-(His)_6_-tagged ribosomes from *E. coli* JE28 strain were purified as described previously (Ederth et al., 2009). An overnight culture of *E. coli* JE28 was used to inoculate 1 liter LB-medium supplemented with 50 μg/ml kanamycin. The cells were grown to an OD_600_ of 1.0 at 37°C. After that, the culture was kept at room temperature for 1 h before shifting it to 4°C for another hour to produce run-off ribosomes. The cells were harvested by centrifugation at 4,000 rpm for 30 min. For purification, the cell pellet was resuspended in lysis buffer (20 mM Tris-HCl pH 7.6, 10 mM MgCl_2_, 150 mM KCl, 30 mM NH_4_Cl) with 0.5 mg/ml lysozyme, 10 μg/ml DNAse I and lysed using a BeadBeater (BioSpec). The lysate was then clarified by centrifugation and passed over a Ni^2+^-chelating column. The ribosomes were eluted with 150 mM imidazole and then dialyzed overnight against lysis buffer. The ribosomes were then pelletized by ultracentrifugation and resuspended in 500 mM NH_4_Cl, 50 mM Tris-acetate, and 25 mM Mg-acetate resulting a final concentration of 10-20 mg/ml. pH was adjusted to 7.2. All buffers were supplemented with complete protease inhibitor cocktail (Roche).

### Labeling of ribosomes after purification

Nonspesific labeling of ribosomes was obtained as described previously (Blanchard, 2004). For this, the purified ribosome complex was incubated with Atto-633-NHS ester in 20 mM Tris-HCl, 10 mM MgCl_2_, 150 mM KCl, 5 mM NH_4_Cl for 30 min at 37°C. Labeled ribosomes were then separated from unbound dye by ultracentrifugation for 3 h at 40,000 rpm and 4°C using a Beckman Coulter ultracentrifuge (Rotor Type 90 Ti). The pellet containing the ribosomes was washed four times with 140 μl 150 mM KCl, 50 mM K-HEPES and then resuspended in the same buffer. pH was always 7.5.

### Single molecule microscopy measurements

Single-molecule fluorescence microscopy measurements were performed on the inverse microscope Olympus IX83 equipped with Toptica iBeam-Smart lasers for 640 and 488 nm and Andor iXon3 EMCCD camera. The YidC-vesicles were mixed with empty liposomes in a ratio of 1:1000. The empty liposomes were prepared from DOPE/DOPG 7/3 mol. % by extrusion technique using polycarbonate membranes with a pore diameter of 100 nm (Avestin, Canada). Synthetic lipids DOPE and DOPG were used instead of *E. coli* polar lipid extract to avoid lipid autofluorescence.

### Fluorescence correlation spectroscopy (FCS)

Fluorescence correlation spectroscopy (FCS) detected ribosome binding to proteoliposomes and determined the number of proteins per vesicle. In brief, the average residence time *τ*_D_ and the number of labeled ribosomes in the confocal volume were derived from the autocorrelation function G(τ) of the temporal fluorescence signal, which was acquired using a commercial laser scanning microscope equipped with avalanche diodes (LSM 510 META ConfoCor 3, Carl Zeiss, Jena, Germany) equipped with a 40x water immersion objective. The diffusion coefficient D was determined as *ω*^2^/4*τ*_D_, where *ω* is the radius of the focal plane. The residence time *τ*_D_ was obtained from the standard model for one-component free 3D-diffusion (Magde et al., 1974):

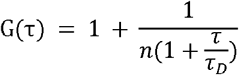

where n is the number of fluorescent particles in the confocal volume.

To determine the number of proteins per vesicle (Hoomann et al., 2013), we first counted the number of proteoliposomes in the confocal volume using the ATTO-488-label on YidC. Second, we compared the number of particles per confocal volume before and after the solubilization of YidC-vesicles with 3 % octyl glucoside (OG). The detergent concentration was chosen to be well above the critical micelle concentration and therefore solubilizes the liposomes. The increase in the number of particles then provides a good estimation of proteins per vesicle. Consequent addition of 1.1 % SDS was performed to examine a possible oligomeric state of YidC in OG micelles.

### Ribosome nascent chains

RNCs of F_0_c were prepared *in vivo* from *E. coli* KC6(DE3) harboring pBAD-F0c(1–46) (a gift from E.O. van der Sluis and Roland Beckmann (Wickles et al., 2014)). To generate RNCs *in vivo*, cells were grown at 37 °C to an OD600 of 0.5 and induced for 1 h with 0.2 % arabinose. Subsequently, the cultures were cooled down on ice, then harvested by centrifugation for 10 minutes in a SLC 6000 rotor at 5500 rpm and 4 °C. The cell pellet was resuspended in 1.5 ml RNC buffer per gram of wet cell pellet (50 mM Tea-acetate, pH 7.5; 150 mM KOAc, 10 mM Mg(OAc)_2_, 1 mM tryptophan, 250 mM sucrose) in the presence of cOmplete EDTA-free protease inhibitor. Cells were lysed by passing them through a French press at 8000 psi. A final concentration of 0.1 % DDM was added. Cell debris was removed at 16,000 rpm and 4 °C for 20 minutes in a SS34 rotor. To separate RNCs from membranes and the cytosolic fraction, the lysate was overlaid on a sucrose cushion (RNC buffer with 750 mM sucrose) in a 1 to 2 ratio and centrifuged in a Ti50.2 rotor for 17 h at 24,000 rpm and 4 °C. The pellet was then resuspended in 4 ml RNC buffer supplemented with 0.1 % DDM by vigorous shaking. After complete resuspension, it was applied onto pre-equilibrated TALON material (2 ml slurry washed once with water, twice with RNC buffer with 0.1 % DDM and once with RNC buffer supplemented with 0.1 % DDM and 10 μg/ml yeast in a propylene column). After 1 hour of binding at 4 °C, the TALON material was washed five times with 10 column volumes of RNC buffer lacking tryptophan. RNCs were eluted by 6 consecutive elution steps of 1 ml RNC elution buffer (50 mM Tea-acetate, pH 7.5, 150 mM KOAc, 10 mM Mg(OAc)2, 150 mM imidazole, 250 mM sucrose, cOmplete EDTA-free protease inhibitor). The eluted fractions were pooled, and RNCs were collected for 2 h in a TLA100.3 rotor at 40,000 rpm and 4 °C. The ribosomal pellet was resuspended in 300 μl buffer (50 mM Tea-acetate, pH 7.5; 150 mM KOAc, 10 mM Mg(OAc)_2_), aliquoted, flash frozen in liquid N_2_ and stored at -80 °C.

### Calculation of channel ion selectivity

The potential, ψ *r*, at which the ionic current through the channel equals zero, was taken from either single-channel analysis or current-voltage ramp recordings. From this so-called reversal potential, the anion (Cl^-^) to cation (K^+^) permeability ratio r (P_Cl_/P_K_) was calculated according to Goldman’s equation for bi-ionic potentials, with ion concentrations K^+^_X_ and Cl^-^_X_ in *cis* and *trans* compartments:

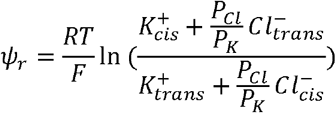

For potassium and chloride concentrations in the two compartments, we account for the osmotic water flow within the unstirred water layers near the membrane, which concentrates the solution on the hypoosmotic side and dilutes it on the hyperosmotic side of the membrane. For such membranes, this effect does not usually exceed 10 %. Therefore we assumed the bulk KCl gradient of 450 to 150 mM to correspond to a 230 mM gradient adjacent to the lipid bilayer.

### Estimation of the pore diameter

A rough estimation of the pore diameter can be obtained from the single-channel conductance using

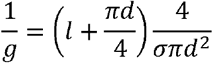

where d, l, and σ are channel diameter, channel length, and conductivity of the solution, respectively (Saparov et al., 2007). σ=3.7 S/m was taken as an average from the conductivities of the two compartments, the channel length was assumed to be l=3 nm.

### Modeling of YidC dimers with AlphaFold

For modeling likely YidC dimer conformations, the *E. coli* YidC sequence (UniProt: P25714) was used as the query_sequence in ColabFold (version 1.5.2) (Mirdita et al., 2022) for GoogleColab, which utilizes AlphaFold2 (Pahari et al., 2019) and AlphaFold2-multimer (Evans et al., 2021) as well as MMseqs2 (Mirdita et al., 2019) and HHsearch (Steinegger et al., 2019) for alignment/template search. Settings were largely kept as default. *AlphaFold2_multimer_v3* was used as the AlphaFold *model_type*. The resulting models were analyzed concerning their plausibility, and model visualizations were generated in PyMOL (Schrödinger).

## Acknowledgments

We thank Dr. Sanyal (Uppsala University, Sweden) for providing the construct for the expression of His-tagged ribosomes and E.O. van der Sluis and R. Beckman (LMU Munich) for the plasmid pBAD-F0c(1–46)

