## Supplementary Information for "YidC from *Escherichia coli* forms an ion-conducting pore upon activation by ribosomes"

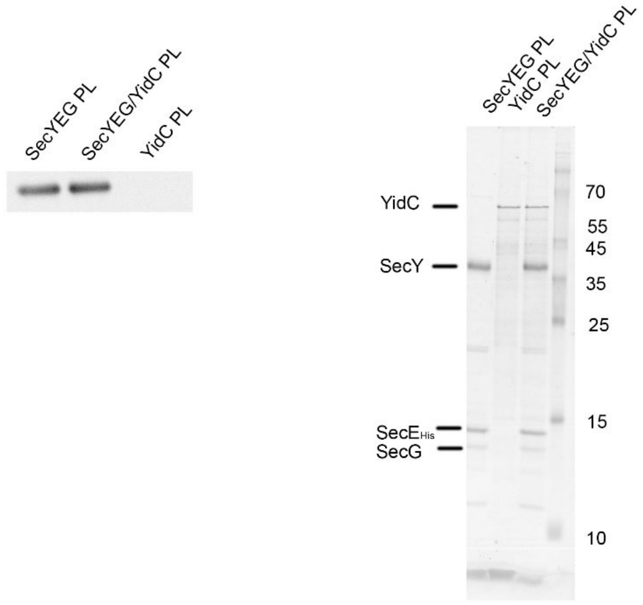


**SFig. 1. Western blotting** demonstrates that purified YidC was not contaminated with SecYEG. The indicated proteoliposomes (25 pmol protein) were separated on SDS-PAGE and after western blotting, the membrane was cut into two pieces. The upper part was decorated with antibodies against YidC and the lower part with antibodies against SecY. YidCΔC corresponds to a YidC derivative lacking the C-terminal 13 amino acids.

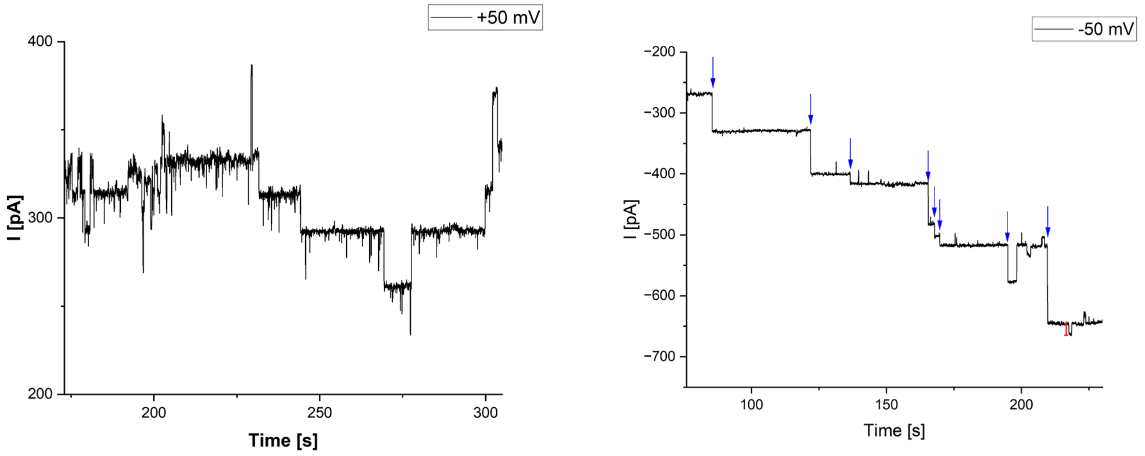


**SFig. 2. Multiple YidC channels activated by ribosomes.** Left: the lipid membrane contained about 10 open YidC channels. Right: fusion events of YidC-proteoliposomes with the planar membrane. The red bar marks the amplitude corresponding to a single YidC channel. At the end of the record, the planar membrane contained more than 20 open YidC channels.

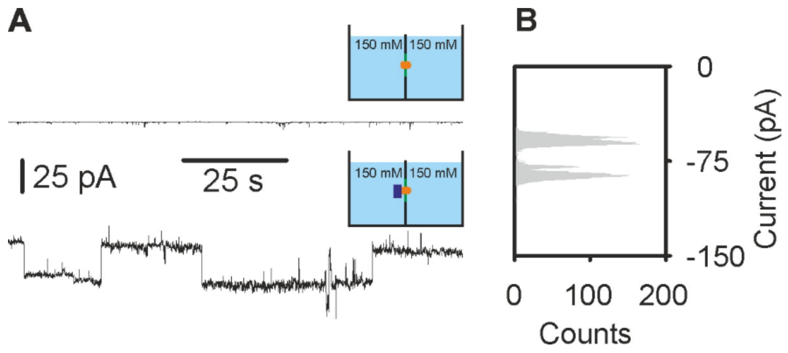


**SFig. 3**. **YidC exhibits channel activity only after ribosome binding. (A)** Traces of current through lipid bilayer containing YidC. The bilayer was folded from monolayers formed on top of a proteoliposome suspension (see *Methods*) in symmetric salt conditions: 150 mM KCl and 50 mM K-HEPES, pH 7.5 in both compartments. Upper trace: negative control. Zero current through the lipid bilayer, containing YidC, before the addition of empty ribosomes. The same was observed when only ribosomes were added to the bare bilayer (data not shown). Lower trace: current through the lipid bilayer showing single channel activity after addition of empty ribosomes. -75 mV were applied across the bilayer. **(B)** Current histogram corresponding to the lower trace in (A). The distance between the two neighbor peaks on the histogram corresponds to the current through a single channel, $I_{SC}$ = -24,7 ± 4 pA or single channel conductivity $g$ = 329 ± 53 pS. Colored schemes show the chamber with two compartments filled with the solution of the indicated ionic strength. The compartments are separated by the lipid bilayer (green), containing YidC (orange) and bound ribosome (purple).

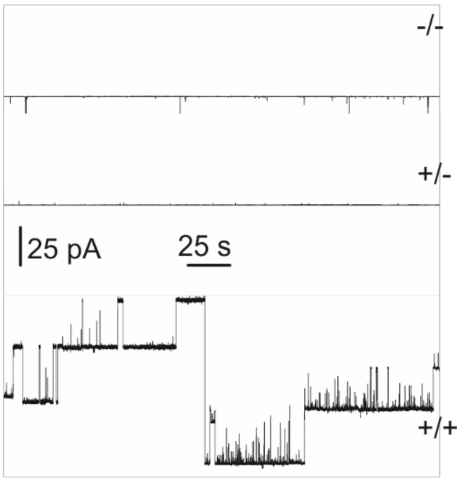


**SFig. 4. Signal peptide is not able to open YidC channels.** Single channel traces shown were recorded at a transmembrane salt gradient of 450 : 150 mM KCl and a transmembrane potential of -50 mV. Both compartments were buffered (pH 7.5) by 50 mM K-HEPES. The upper panel shows no channel activity in the absence of both, the signal peptide of cytochrome O oxidase’s A subunit and the ribosome (-/-). Upon addition of solely signal peptide (+/-), no channel activity could be observed. The further addition of ribosomes (+/+) resulted in channel activity.
